# Quantitative proteomics and phosphoproteomics analyses identify sex-biased protein ontologies of *Schistosoma japonicum*

**DOI:** 10.1101/2025.05.30.656995

**Authors:** Chuantao Fang, Bikash R. Giri, Guofeng Cheng

## Abstract

*Schistosoma japonicum* (*S. japonicum*) is one of the major causative agents of human schistosomiasis in Asia. Identification of differentially expressed proteins (DEPs) between male and female worms could reveal critical signaling pathways that are involved in sexual maturation and egg production. In the study, quantitative proteome and phosphoproteome profiles were obtained from adult male and female worms of *S. japonicum*. A total of 2,710 unique proteins, including 2,055 proteins and 924 phosphorylated proteins, were identified. We identifed 252 (∼12.5%) non-phosphorylated and 209 (11.7%) phosphorylated DEPs between males and females. Combined with the RNA sequencing (RNA-seq) results, 22 non-phosphorylated DEPs exhibited corresponding mRNA-level changes. Next, several identified non-phosphorylated DEPs were further shown to be involved in sex-biased biological processes, including vitellocyte development, oviposition, and parasite mobility by RNA interference, which were also supported by their cellular niches by analyzing previously published single-cell RNA seq data. Furthermore, we annotated 96 kinases of *S. japonicum* based on the obtained phosphorylated proteins, of which CMGC/MAPK, Atypical/RIO are significantly activated in males, while CAMK/CAMKL, AGC/DMPK, and STE/STE7 are activated in females. Finally, the potential drugs targeting these kinases were determined in silico, resulting in the identification of 28 *S. japonicum* kinases as potentially targetable by 30 FDA-approved drugs. Overall, our study provided a collection of evidence-based proteomic and phosphoproteomic resources of *S. japonicum* and identified sex-biased proteins, phosphopeptides, and kinases, which could serve as potentially effective targets for developing novel interventions against schistosomiasis.

## Introduction

Schistosomes are blood-dwelling helminth parasites that can infect humans and certain mammals worldwide [1]. Schistosomiasis caused by schistosome infections afflicted approximately 230 million individuals in 78 nations, representing a significant global health problem [2]. It is estimated to cause approximately 200,000 deaths annually, contributing 3.3 million disability-adjusted life years (DALYs) to the global disease burden [3-5]. In Asia, *Schistosoma japonicum* (*S. japonicum*) infection was frequently found in China, Philippines, and a small area of Indonesia [6]. Currently, praziquantel is the only drug of choice for schistosomiasis treatment, although it’s ineffective for juvenile worms, of which the long-term usage, unfortunately, was challenged due to the concerns of drug resistance [7, 8]. In schistosomiasis pathogenesis, the primary disease manifestations and transmission dynamics are driven by egg deposition from sexually mature worm pairs. This life cycle dependency highlights multiple potential therapeutic targets. However, translational exploitation of these opportunities has been hindered by limited understanding of the molecular mechanisms regulating parasite development and oviposition.

During the complex life cycle, schistosomes undergo multiple morphological changes, and sexual maturation starts with a continuous pairing of males with females to ensure reproductive development [9, 10]. Previous studies have demonstrated that both physical and chemical intersexual interactions critically regulate reproductive development and egg production in schistosomes [11]. Consequently, elucidating the molecular mechanisms underlying sexually dimorphic developmental processes in schistosomes has emerged as a critical research priority with important therapeutic implications. In 2022, proteomic profiles of unisexual- and bisexual-infected *S. japonicum* male worms were identified to show 674 differentially expressed proteins (DEPs) across multiple developmental stages, suggesting dynamic sex-dependent molecular regulation [12]. Moreover, several studies have established protein phosphorylation as a crucial post-translational modification that orchestrates signaling networks to regulate fundamental biological processes in parasites, including growth and reproductive development [13, 14]. For instance, protein phosphorylation has been demonstrated to play an essential regulatory role in the cercariae-to-schistosomula transformation, a critical step for host infectivity [15]. Similarly, Ressurreicao and coworkers demonstrated that heightened ERK and PKC activation within the cephalic ganglia of praziquantel-treated specimens correlates with tegumental and excretory system modifications, contributing to *S. mansoni* homeostasis [14]. In *S. japonicum*, Luo and coworkers identified 127 distinct phosphorylation sites in 92 proteins, of which 30 were phosphorylated, including signaling molecules like 14-3-3 and HSP90 [16]. These studies highlight the fundamental importance of protein phosphorylation in schistosome biology; however, whether there is a divergence in the phosphoproteome between male and female worms remains unexplored.

In the present study, we employed an integrated iTRAQ quantitative proteomic and phosphoproteomic methodology to pursue two primary research goals: 1) establish a systematic protein expression atlas of *S. japonicum* across sexes, and 2) comprehensively delineate sexually dimorphic protein and phosphoprotein expression profiles between adult male and female worms. In parallel, functional experiments were conducted to validate the biological functions of prioritized molecules. Kinase activity profiling was conducted using a tailored GSVA algorithm, incorporating STRING-derived protein-protein interaction networks as topological frameworks. This multi-omics approach yielded three key outcomes: (1) the first quantitative proteomic/phosphoproteomic resource for *S. japonicum*, (2) identification of sex-biased differentially expressed proteins (DEPs) and phosphoproteins (DEPPs), and (3) characterization of sexually dimorphic kinase activation states.

## Results

### Overview of proteome and phosphoproteome profiles

Adult male and female *S. japonicum* at 35 days post-infection (dpi) were profiled using integrated iTRAQ-based quantitative proteomic (iBQP) and phosphoproteomic (iBQPP) approaches as outlined in the workflow (**Figure 1A**). Briefly, protein lysates from adult male and female worms were independently prepared and tryptically digested to generate peptides. The resultant peptides underwent 8-plex iTRAQ isotopic labeling, with subsequent phosphopeptide enrichment via titanium dioxide (TiO_2_) affinity chromatography. Triplicate biological replicates were carried out by Q-Exactive liquid chromatography-tandem mass spectrometry (LC-MS/MS) system following quality control validation.

**Figure 1.**
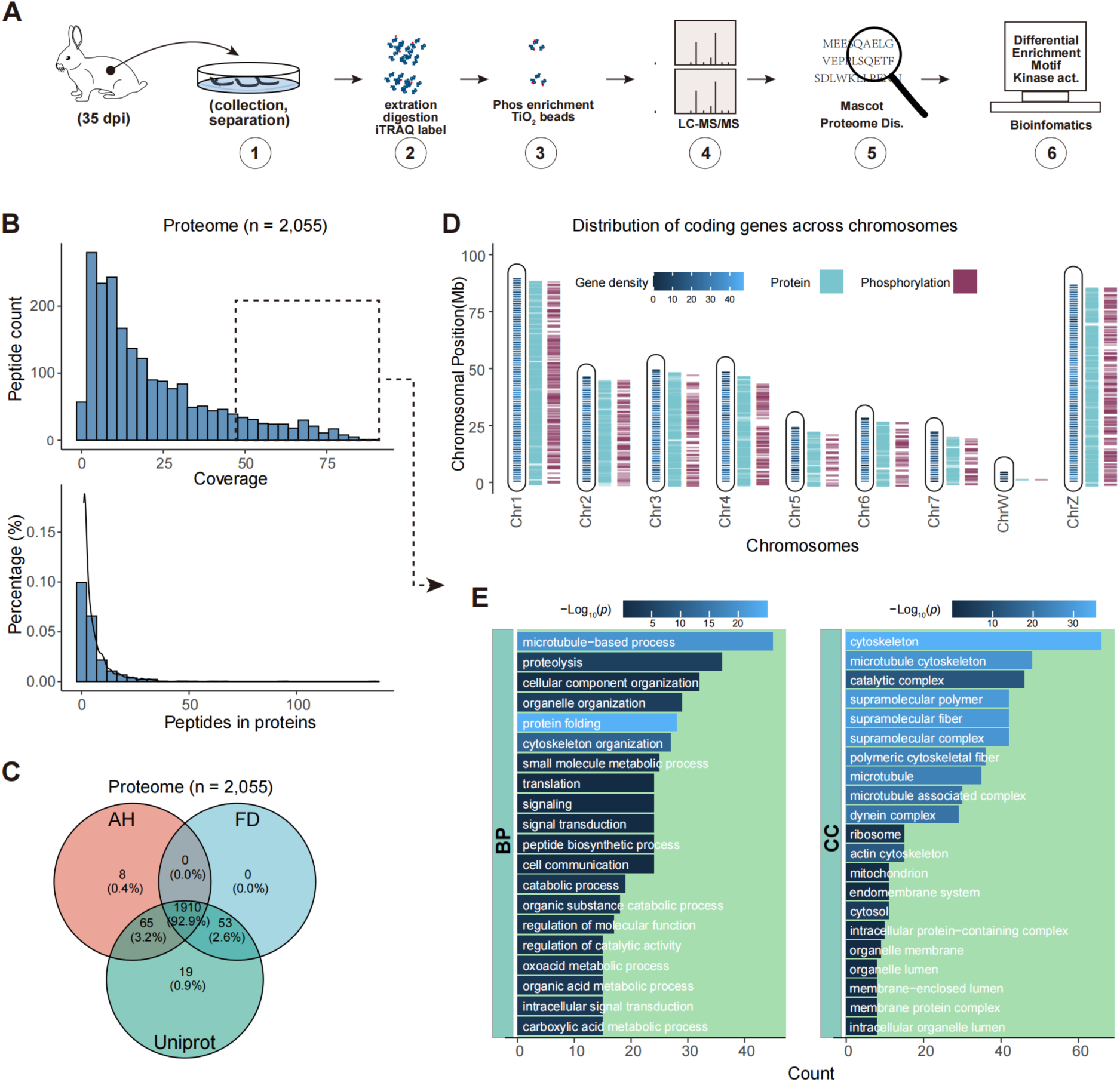
Overview of quantitative proteome and phosphoproteome profiles. **A**. Workflow (including animal model construct, worm collection/separation, protein sample preparation, LC-MS/MS, and bioinformatics) of the current study. Dpi, days post-infection; iTRAQ, Isobaric Tags for Relative and Absolute Quantitation. **B**. Distribution of unique peptide coverage (Top) and total peptide (Bottom) of 2,055 proteins identified. Proteins with a high coverage ratio (>50%, in a dashed line rectangle) were subjected to functional enrichment analyses in (E). **C**. Venn plot indicated the relationship between the proteomic data with 3 widely used protein annotations of *S. japonicum*. AH, Anhui version of PRJEA34885 (https://parasite.wormbase.org/Schistosoma_japonicum_prjea34885/Info/Index/); FD, Fudan version of PRJNA520774 (https://parasite.wormbase.org/Schistosoma_japonicum_prjna520774/Info/Index/); and UniProt, retrieved by the keyword of taxonomy_id:6182. **D**. Overview of protein-coding gene (identified by proteomics and phosphoproteomics) locus on 9 chromosomes of *S. japonicum*. The chromosomes were defined by the Genome assembly ASM2521551v1 of *S. japonicum*. **E**. Functional enrichment analyses of highly expressed proteins in the proteome using Gene Ontology (GO) gene sets. BP (biological processes) and CC (cellular component) are defined by the GO database. The top 20 enriched terms were presented.

According to the Mascot program searching, we identified 317,107 raw spectra in total, which were matched to an average of 37,690 spectra and 10,046 high-scoring unique peptides after peptide retrieval using the *S. japonicum* proteome database (Table S1). This resulted in the identification of 2,055 non-redundant proteins with 21.6% sequence coverage (Figure 1B; Tables S1 and S2). Database annotation revealed 99.6% (2,047/2,055) of proteins matched UniProtKB entries, while 0.4% represented gene products from previous genome annotations (ASM15177v1 genome assembly) (Figure 1C). Chromosomal mapping demonstrated non-biased protein distribution across all nine chromosomes, exhibiting congruence with genomic gene density patterns (Figure 1D).

To reflect the most prominent biological characteristics of adult worms from identified proteins, we performed Gene Ontology (GO) enrichment analyses of high-abundance proteins (with sequence coverage >50%) to reveal significant associations with: 1) biological processes: microtubule-based motility (GO:0007018), proteolytic regulation (GO:0006508), organelle reorganization (GO:0006996), and chaperone-mediated folding (GO:0061077); 2) cellular components: cytoskeletal architecture (GO:0005856), catalytic complexes (GO:0098783), and microtubular networks (GO:0005874); 3) molecular functions: GTPase activity (GO:0003924), ATP-dependent catalytic activity (GO:0140657), and ATP hydrolysis-coupled transport (GO:0043461) (Figure 1E, Figure S1A). Consistently, Kyoto Encyclopedia of Genes and Genomes (KEGG) pathway analyses corroborated these findings, showing enrichment in Salmonella infection signaling (sja05132), motor protein interactions (sja04810), and tight junction regulation (sja04530) (Figure S1A).

### Quantitative description and functional enrichment analyses of differentially expressed proteins (DEPs) between males and females

To systematically investigate sex-biased proteomic profiles in *S. japonicum*, we conducted principal component analyses (PCA) on global protein expression data, revealing that primary variance (PC1, 55.3%) predominantly originated from sexual dimorphism (Figure S2A and B). This sexual divergence was corroborated by clustering analyses of protein expression (or proteomic profile), with clear segregation between male and female worms in the heatmap visualization **(Figure 2A**). Integration of our proteomic dataset with transcriptomic data of the same samples revealed a considerable concordance between protein abundance and corresponding mRNA levels (Figure S2C), and RNA levels can explain about 30% variance in protein levels (33% in male and 24% in female). Meanwhile, the comparative analyses demonstrated minimal overlap between differentially expressed proteins (DEPs, n = 252) and differentially expressed genes (DEGs), with 22 genes (such as EWB00-000538, EWB00-007550, EWB00-004736, EWB00-003121, EWB00-001015, EWB00-009171, etc) exhibiting concurrent differential expression at both levels (Figure 2B, S2D and S2E; Table S3). Furthermore, protein-protein interaction network analyses of DEPs identified three functionally significant modules: cytoskeletal remodeling complexes, vacuolar ATPase assemblies, and signal transduction cascades (Figure 2C).

**Figure 2.**
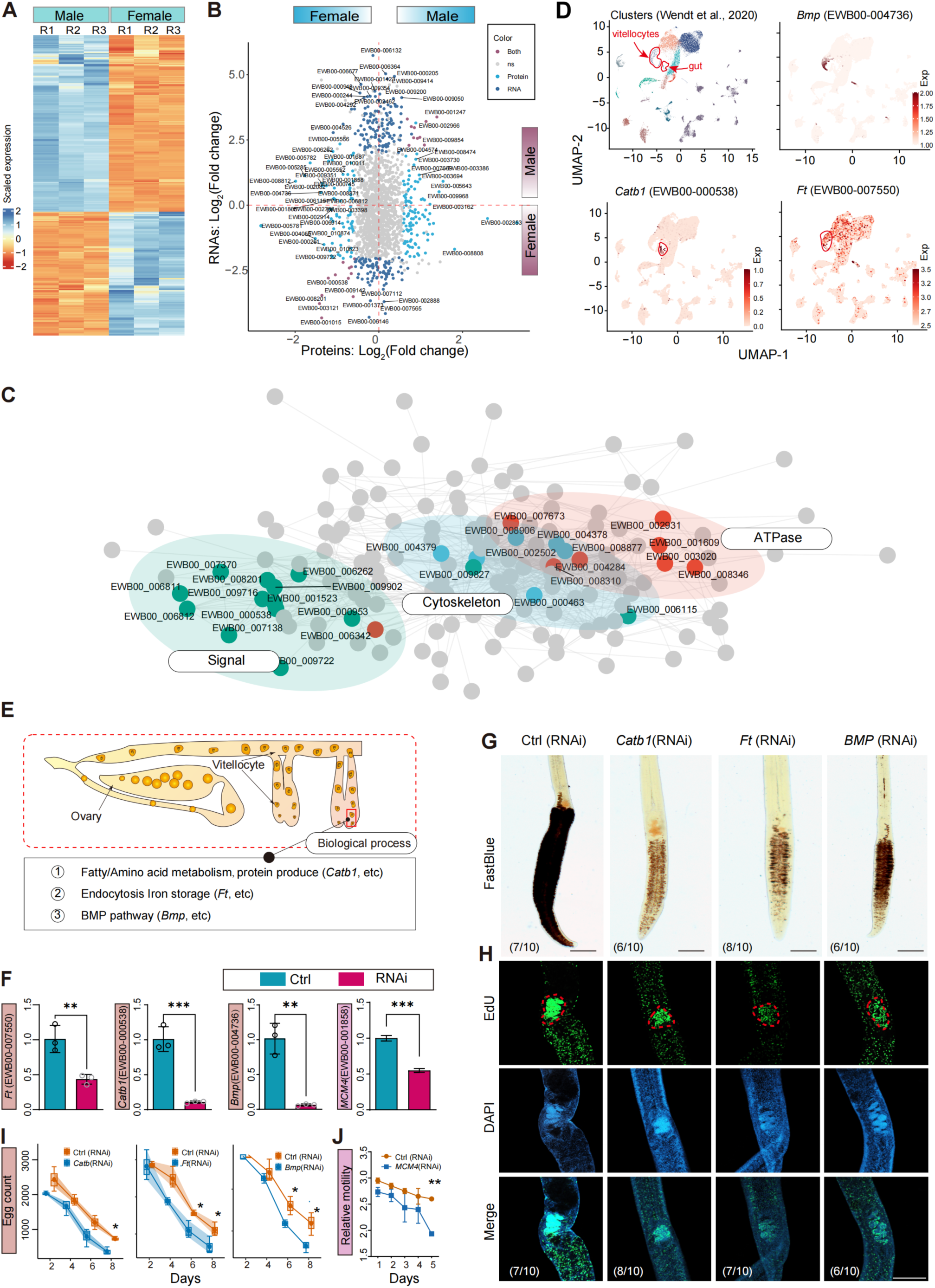
Differentially expressed proteins between male and female worms. **A**. Heatmap showing differentially expressed proteins between male and female worms in three biological replicates. The cutoff of fold-change>1.5 and adjusted p*<*0.05 was used. exp, expression. **B**. The overlap between differentially expressed genes (in RNA-seq) and proteins (in the proteome of the current study). Both are differentially expressed in protein and mRNA levels; Protein/RNA is only differentially expressed in protein/mRNA levels. **C**. Protein-protein interaction of identified differentially expressed proteins. STRING R package was employed for the analyses. **D**. Annotation of selected proteins in cell clusters from a scRNA-seq dataset of schistosomes. *S. japonicum Catb1, Ft,* and *Bmp* expression in two-dimensional Uniform Manifold Approximation and Projection (UMAP) space of scRNA sequencing data of *S. mansoni*. Exp, expression in a scRNA sequencing dataset. **E**. The model showing the possible mechanisms underlying vitellocyte development. **F.** Real-time PCR determined the mRNA level of the indicated genes. **p<0.01, and ***p<0.001 by student’s *t*-test. **G-H**. FastBlue BB analyses (G) of morphological vitellarium and EdU staining (H) in *Catb1*/*Ft/Bmp* inhibited females. Worms targeted the *Catb1*/*Ft/Bmp* via RNAi were used. n = 10, scale bar, 100 μm. RNAi, RNA interference. Numbers in the corner indicated that the parasites showed similar alternation in the vitellarium; a red circle showing the ovary. **I**. Egg count of the females treated with *Catb1/Ft/Bmp* dsRNA after 2 d, 4 d, 6 d, and 8 d. The same worms in (F) was used. *p*<*0.05 by one-way ANOVA analyses. **J.** Worm mobility showed the functional alteration of worms after knocking down sex-specific coding genes. **p<0.01 by ANOVA.

Because the pathological injuries to the host of *S. japonicum* are mainly caused by egg deposition in host organs, especially in the livers, we mainly focused on the vitellarium development of female males, which is responsible for egg formation and production. Therefore, we integrated our results with the published single-cell RNA (scRNA) sequencing data of *S. mansoni* [17]. Interestingly, we found that more than 15 DEGs were female-enriched (especially vitellocyte-enriched) genes, including EWB00-000538, EWB00-004736, EWB00-001805, EWB00-007550, EWB00-005782, etc (Figure 2D and S2F).

Protein-protein interaction analyses of these proteins using the STRING database suggested their functions related to the followings: 1) biological synthesis, 2) substance transport, and 3) BMP signaling (Figure 2E). In addition, these processes may be necessary for the vitellocyte development, which is naturally a cell proliferation process that requires biological factors and cellular signaling. Therefore, we selected 3 genes, *Catb1* (EWB00-000538), *Ft* (EWB00-007550), and *Bmp* (EWB00-004736) to determine their biological functions in female worms (Figure 2F-I). Interestingly, we observed that knockdown of these genes by RNAi can influence the vitellocyte development and ovarian cell proliferation activity (Figure 2G and H). Moreover, the number of produced eggs also significantly decreased in the inhibited females (Figure 2I). Also, mobility of males is linked to mating stability and then the following sexual maturation of schistosomes [18]. We identified two male-enriched genes (*Mcm4* and *Agp*) that were associated with male mobility (Figure 2F and J, Figure S3B and C). Collectively, these results demonstrate that the DEPs obtained from proteomic profiling provide important insights for understanding the mechanisms of sex maturation and egg production in *S. japonicum*.

### Differentially expressed phosphorylated proteins between males and females

Phosphorylation, as the most prevalent post-translational modification in eukaryotic species, serves as a critical regulatory mechanism across diverse biological pathways [19, 20]. Despite its evolutionary conservation in higher organisms, the phosphoproteomic landscape and its functional implications remain poorly characterized in *S. japonicum*. To address this knowledge gap, we performed a comparative phosphoproteomic analysis between male and female worms. Initial quality assessments revealed statistically comparable parameters in intersex comparative phosphoproteomic datasets, including total peptide counts, peptide length distributions, and quantitative intensity profiles (Figure S4A-C).

Through integrated annotation using the *S. japonicum* proteome database (Anhui, AH, and Fudan, FD version) supplemented with UniProt, we identified 869 phosphoproteins with high confidence (**Figure 3A**; Table S5). Notably, 24.7% (214/869) of total phosphoproteins were identified in the proteomic datasets mentioned above (Figure 3A). Site-specific analyses demonstrated rigorous localization reliability, with>90% of phosphorylation sites (both singly- and multiply-phosphorylated peptides) achieving a posterior probability threshold>0.8 (Figure 3B). Among these high-confidence sites, most peptides possessed 1-2 serine or threonine modifications (Figure 3C). PCA analyses of all phosphoproteins revealed that sexual dimorphism explained most of the phosphoproteomic variance (PC1, 78.2%) (Figure 3D). Hierarchical clustering of phosphoprotein expression levels further confirmed significant intersex divergence (Figure 3E; **Tables 1, 2** and S6). These findings collectively implicate protein phosphorylation as a potential molecular determinant in sexually dimorphic physiological processes in *S. japonicum*.

**Figure 3.**
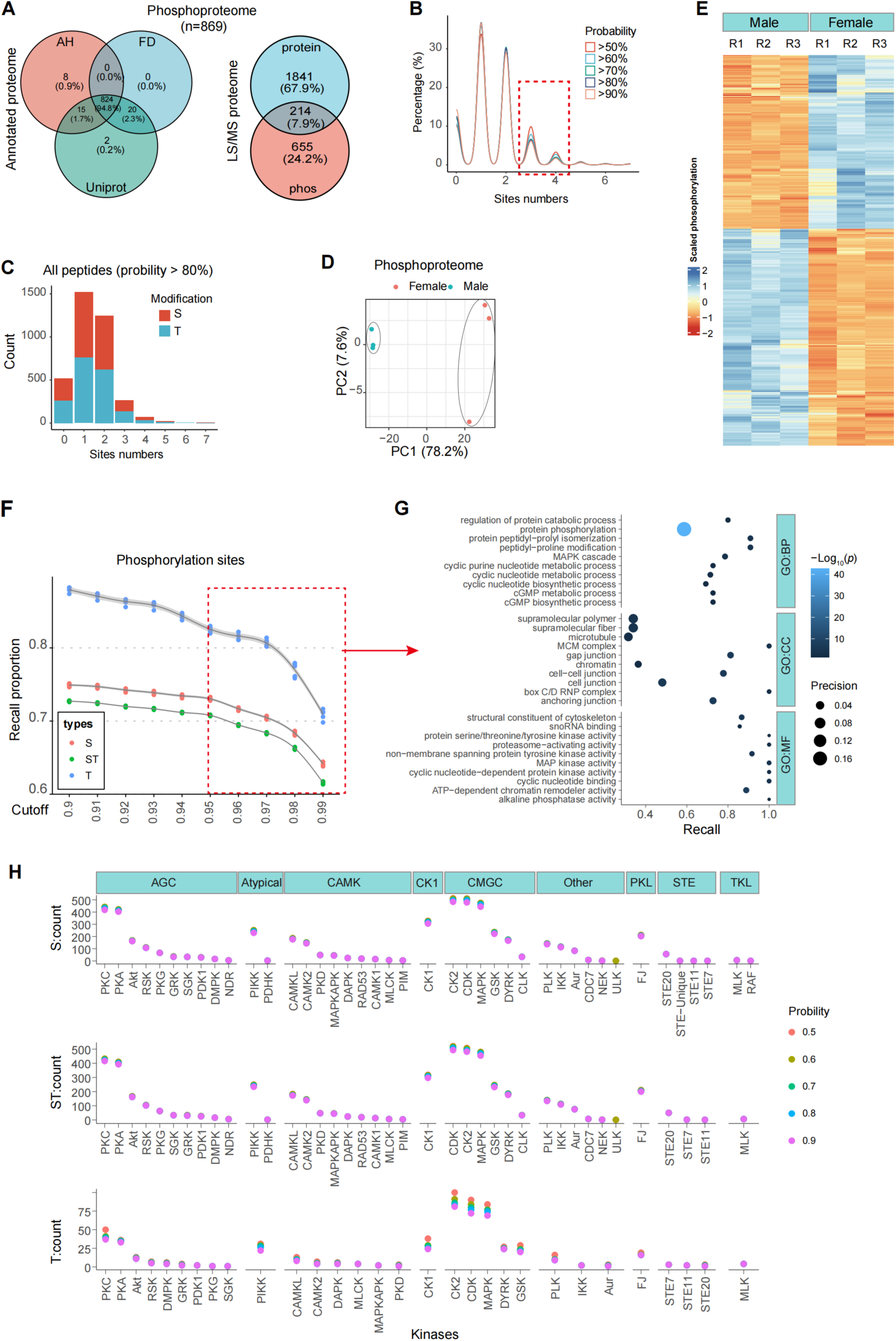
Differentially expressed phosphorylated proteins between male and female worms. **A**. Venn plot indicated the relationship between the proteomic data with 3 widely used protein annotations of *S. japonicum* and the mentioned proteome data. AH, Anhui version of PRJEA34885 (https://parasite.wormbase.org/Schistosoma_japonicum_prjea34885/Info/Index/); FD, Fudan version of PRJNA520774 (https://parasite.wormbase.org/Schistosoma_japonicum_prjna520774/Info/Index/); and Uniprot, retrieved by the keyword of taxonomy_id:6182. **B**. Distribution of the number of peptide-modified sites with the indicated probability. **C**. The number of peptide S/T modifications. **D**. Principal component analyses of phosphoproteomic data. **E**. Heatmap showing differentially expressed phosphorylated proteins between male and female worms in three biological replicates. The cutoff of fold-change>1.5 and adjusted *p<*0.05 was used. **F**. The recall proportion of identified phosphorylation sites by KinasePhos software, kinase-specific phosphorylation site prediction tools (https://awi.cuhk.edu.cn/KinasePhos/About%20KinasePhos.html). Probability, the probability of being a real modification. **G**. Functional enrichment of highly probable phosphorylated proteins by gene ontology-defined gene sets. The top 10 enriched terms were presented. **H**.The total number of predicted phosphorylation sites (with probability>0.5) of the indicated kinases. Kinases of S/T/ST modification were predicted by GPS 6.0.

**Table 1.**
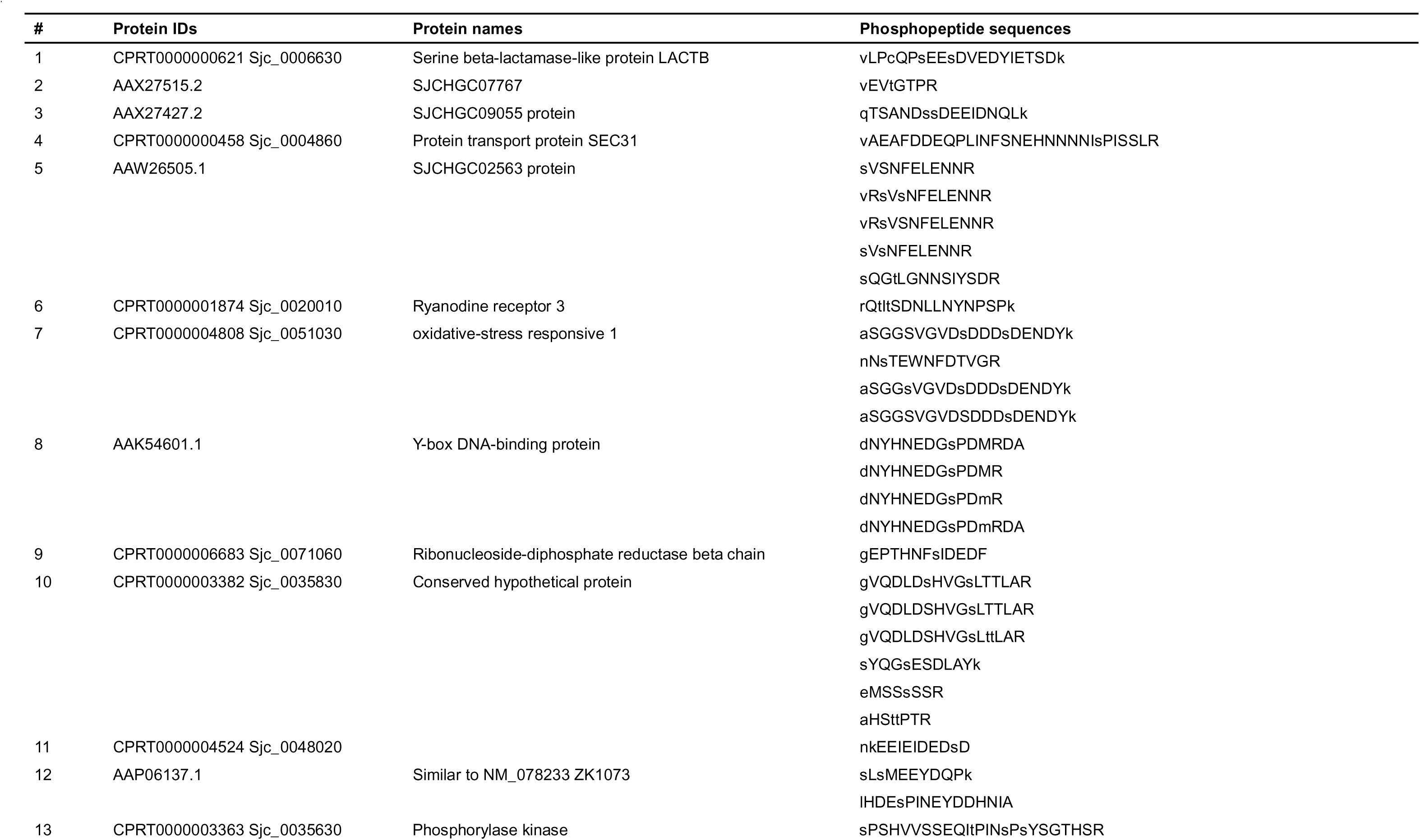
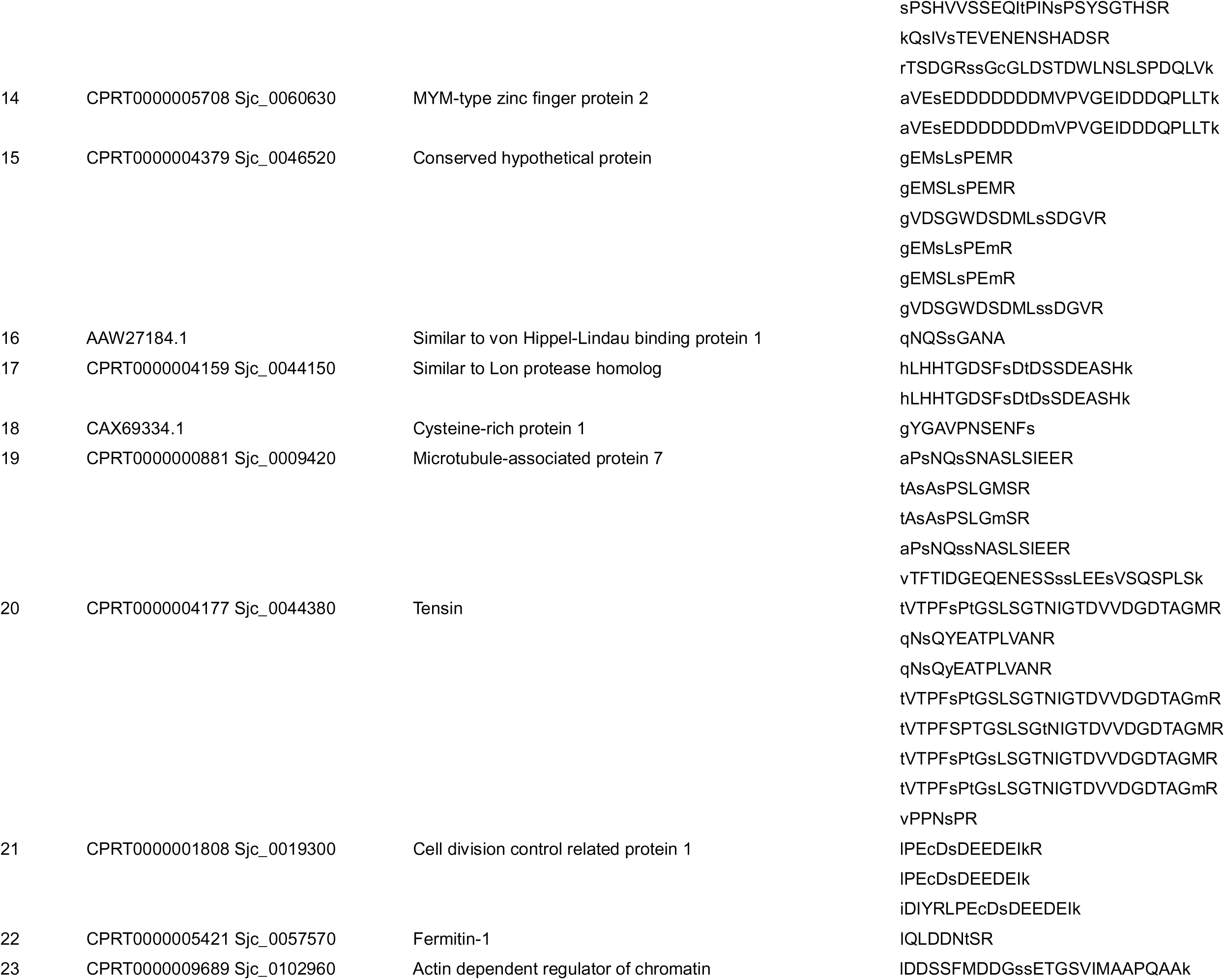
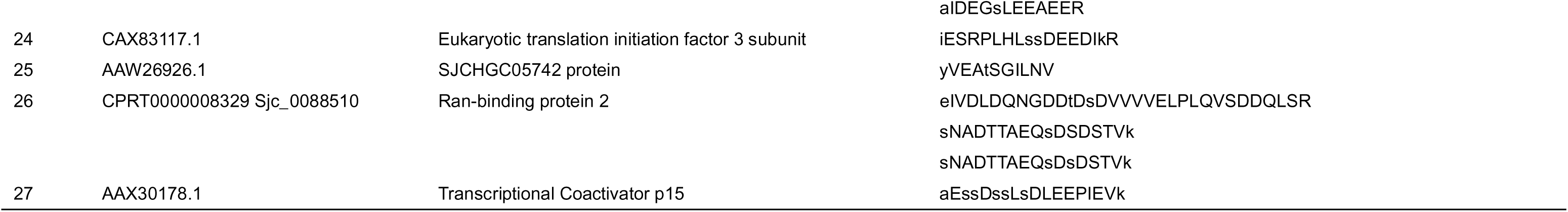
Female enriched phosphorylated proteins in *Schistosoma japonicum*.

**Table 2.**
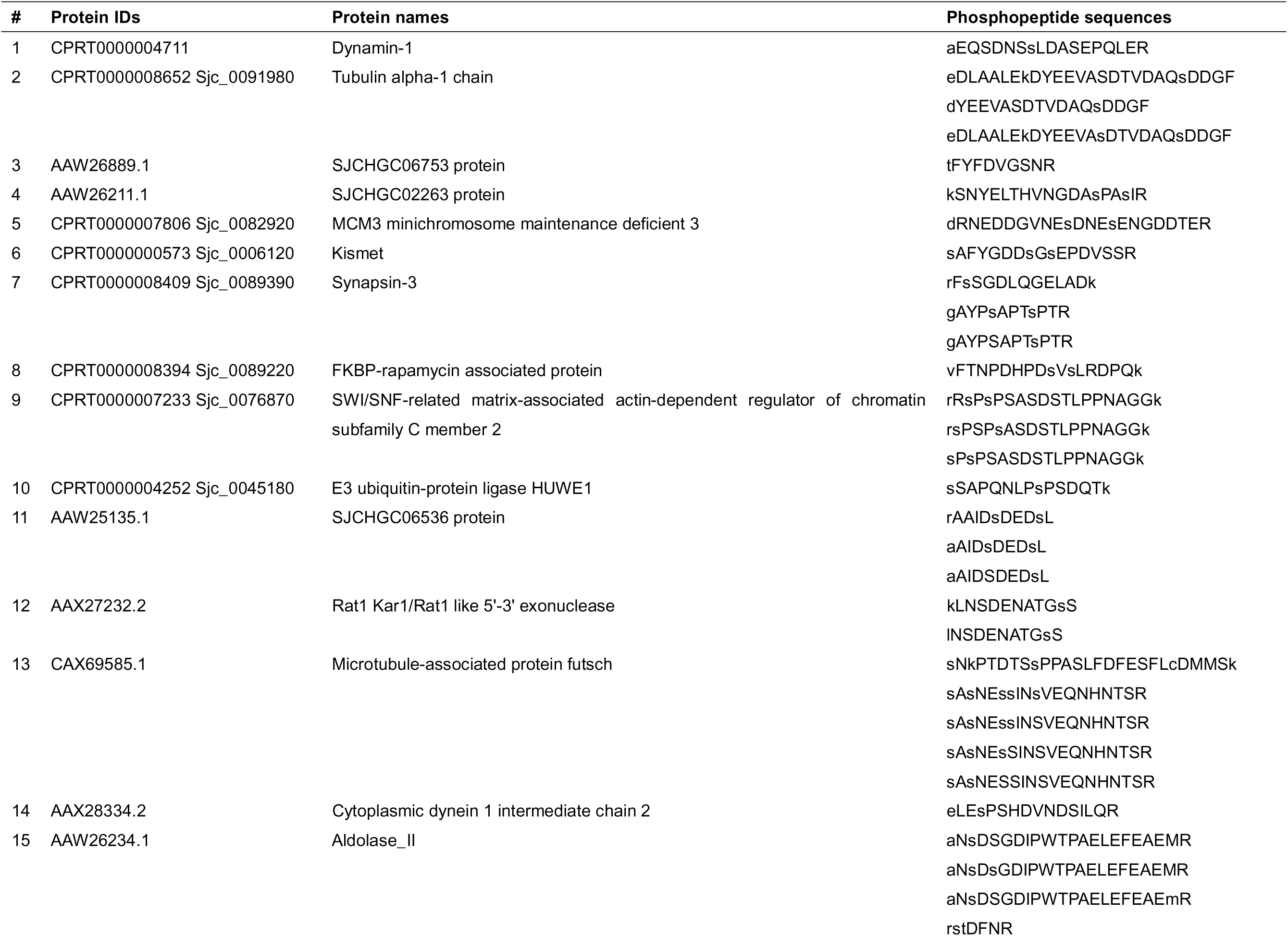
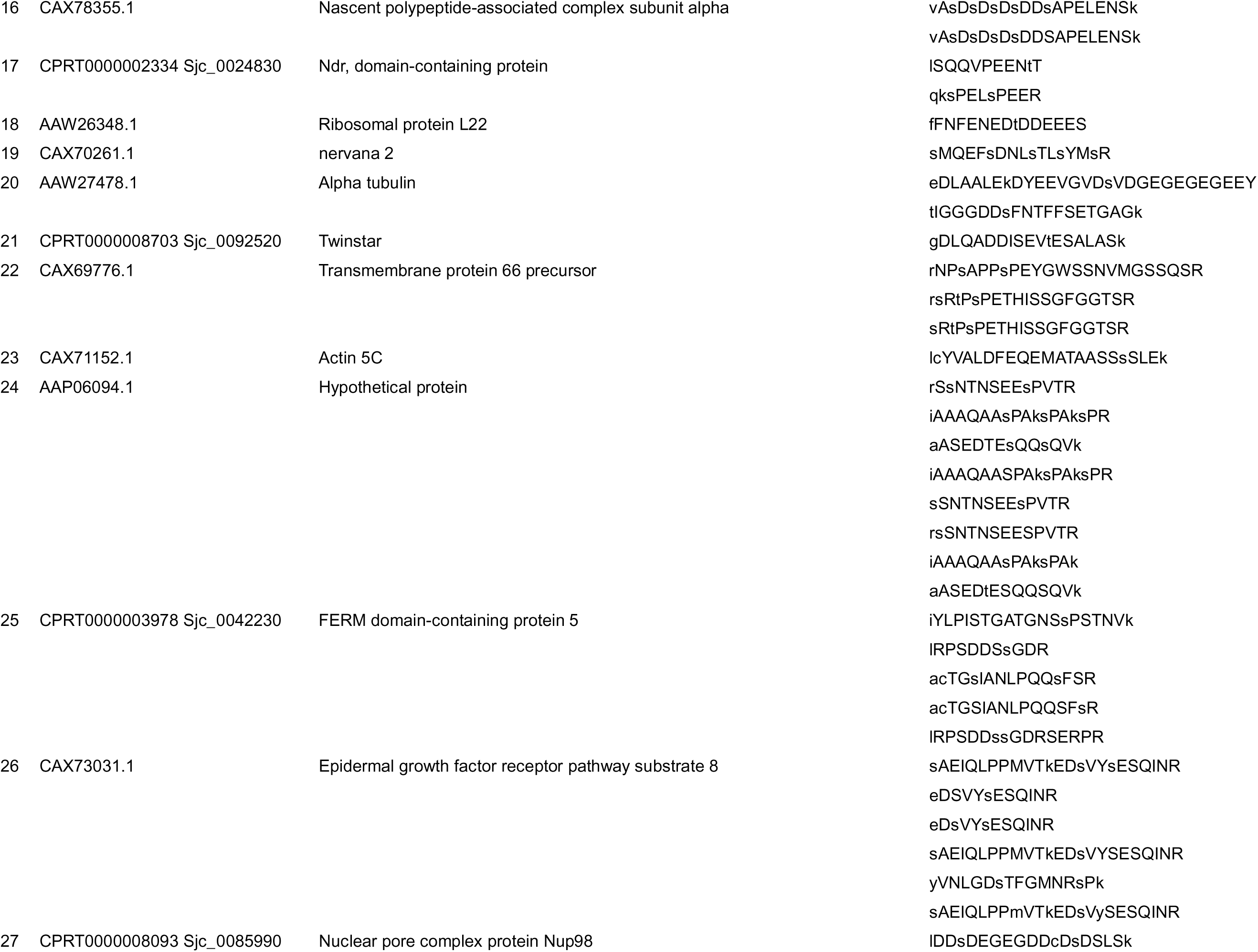
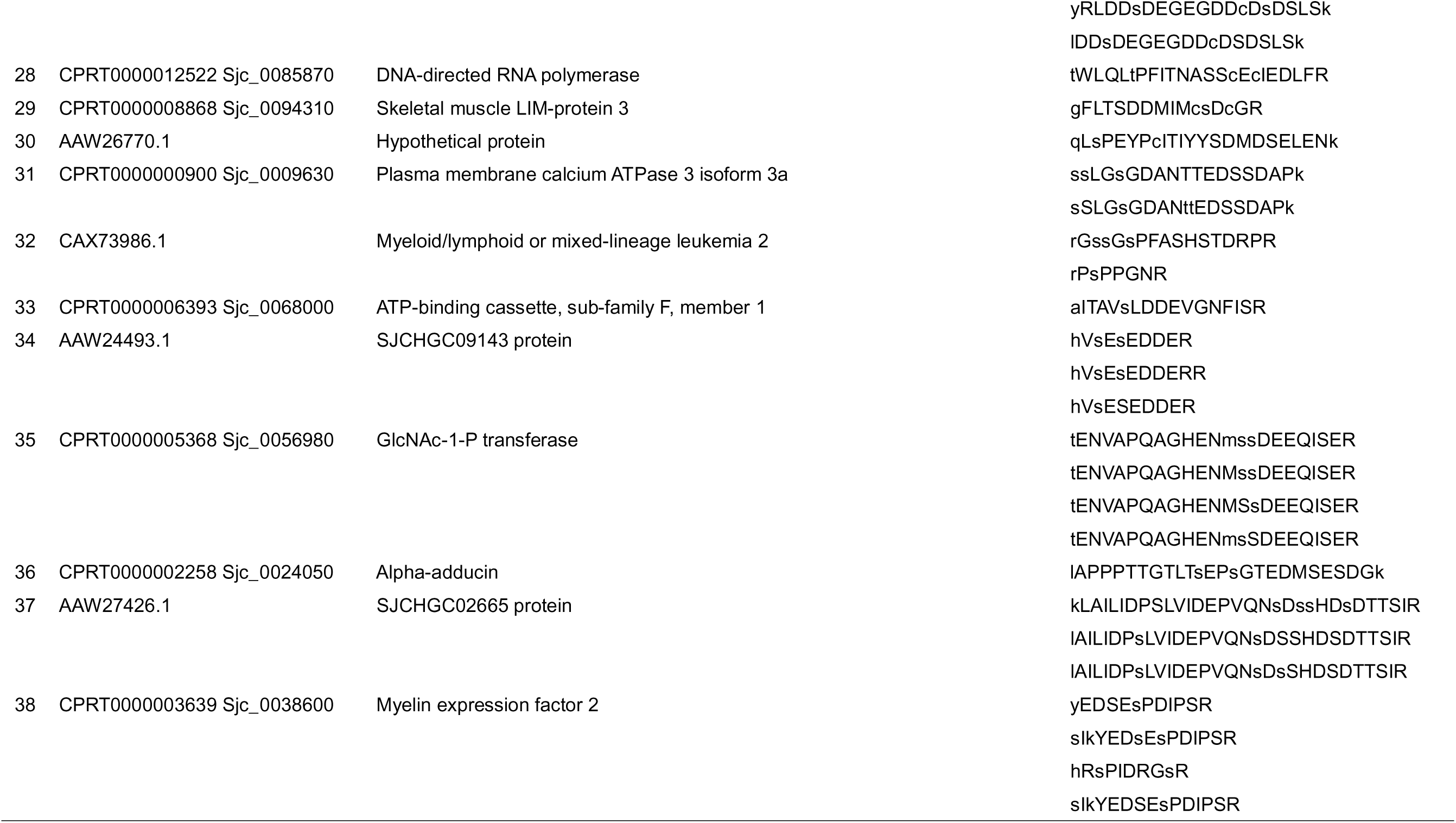
Male enriched phosphorylated proteins in *Schistosoma japonicum*.

To elucidate the biological significance of protein phosphorylation in *S. japonicum*, we performed systematic kinase prediction for serine/threonine/serine-threonine (S/T/ST) phosphorylation sites using KinasePhos3.0, a comprehensive computational tool employing profile hidden Markov models (HMMs) for kinase-specific phosphorylation site prediction [21]. Applying a stringent probability threshold (> 0.95) of being a real modification, we identified 1,428 serine, 1,501 serine-threonine, and 222 threonine phosphorylation sites, respectively (Figure 3F). Functional enrichment analyses revealed these (both detected and predicted) phosphoproteins predominantly participate in 1) biological processes: protein catabolic regulation, phosphorylation cascades, and MAPK signaling pathways; 2) cellular components: supramolecular polymers and microtubule networks; 3) molecular functions: cytoskeletal structural maintenance, serine/threonine/tyrosine kinase activity, and MAP kinase activation (Figure 3G).

Leveraging the S/T/ST phosphorylation sites, subsequent kinome prediction utilizing GPS 6.0 software [22], an enhanced algorithm for kinase-specific phosphorylation site identification, demonstrated preferential modification by three major kinase families: AGC (protein kinases C and A), CAMK (calcium/calmodulin-dependent kinases including CAMKL and CAMK2), and CMGC (cyclin-dependent kinases and MAP kinases). Quantitative distribution across kinase families was quite balanced (Figure S4D). Motif analyses via MEME Suite identified several conserved phosphorylation motifs, notably STE20, CDK, MAPK, and CAMKL signatures (Figure S5). Collectively, these findings suggest a complex, phosphorylation-driven regulatory network in *S. japonicum*, which may be involved in developmental processes and cellular architecture homeostasis via sex-biased kinase families.

### Differential kinase activities between males and females

To comprehensively investigate kinase signaling pathways governing *S. japonicum* development, we developed a computational framework for kinase activity quantification through quantitative phosphorylated substrate profiling. The methodological workflow (Figure 4A) comprises three principal steps: 1) database curation of 29,803 *S. japonicum* protein sequences from UniProt, followed by homology-based prediction of 96 conserved kinases through Diamond [23] alignment (identity threshold>30%) against *Caenorhabditis elegans* kinase orthologs; 2) construction of kinase-substrate interaction networks through the integration of functional and physical protein-protein interaction data; 3) quantitative kinase activity inference via GSVA (Gene Set Variation analyses) [24] applied to phosphoproteome profiles mapped against curated substrate gene sets.

**Figure 4.**
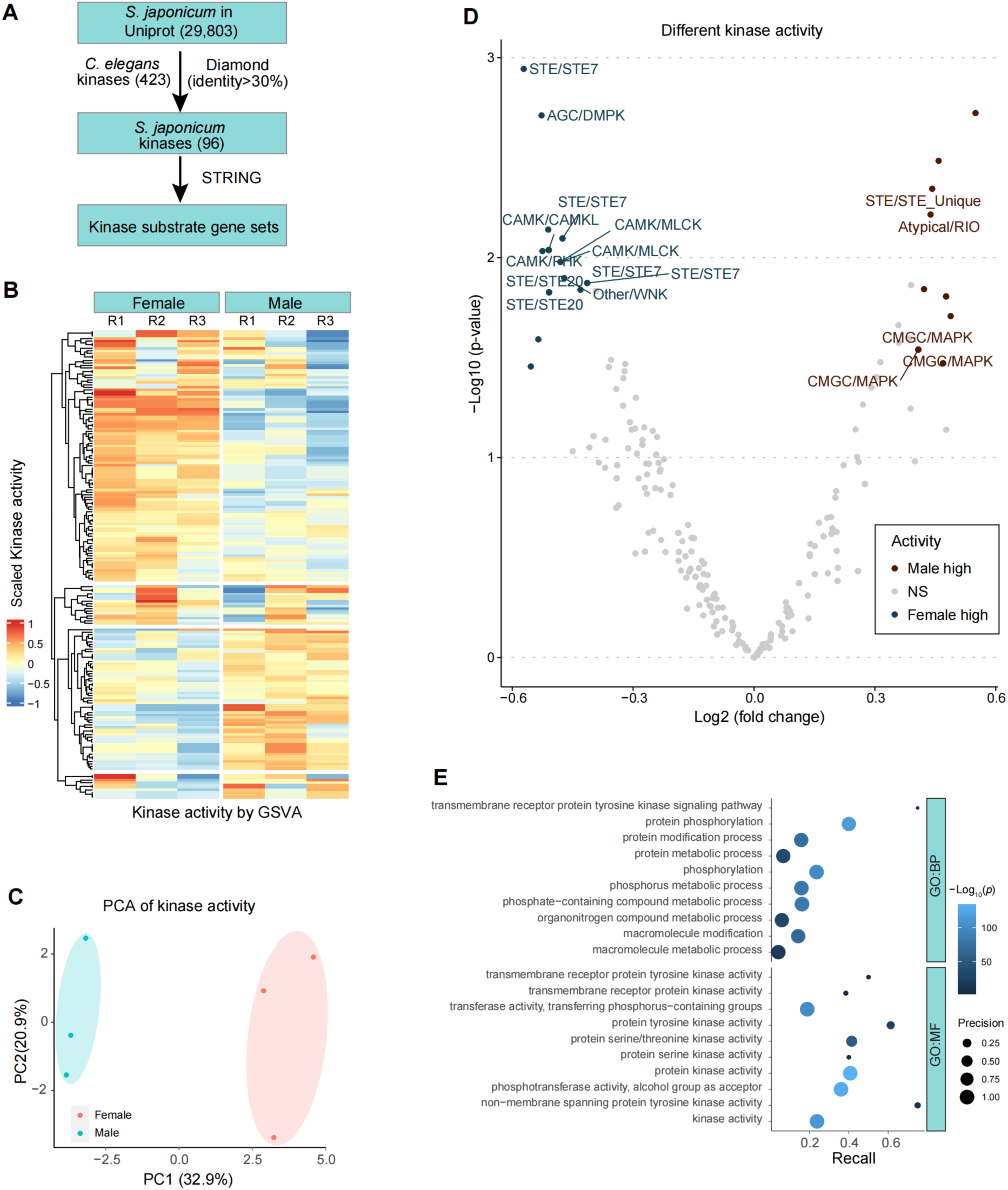
Differentially expressed kinases between males and females. **A**. Flowchart for predicting *S. japonicum* kinases and their interactors. *S. japonicum* kinases were predicted based on protein sequence homology (identity >30%) with *C. elegans* kinases (423). Kinase substrate gene sets were defined by STRING online tools. **B**. Heatmap of kinase activity calculated by the GSVA algorithm. Kinase activity was defined by the enrichment score of kinase substrates in A. **C**. Principal component analyses of kinase activity of female and male worms. **D**. Volcano plot showing the different kinase activities between male and female worms. **E**. GO enrichment analyses of kinases with different activity between female and male worms. The top 10 enriched terms were presented.

Differential analyses of kinase activity revealed significant sexual dimorphism in kinase activity profiles, with pronounced differential activation patterns between male and female worms (Figure 4B and C). Sex-biased kinase enrichment analyses demonstrated male-biased activation of CMGC/MAPK, Atypical/RIO, and STE/STE Unique kinase families, contrasting with female-prevalent activity in CAMK/CAMKL, AGC/DMPK, and STE/STE7 clusters (Figure 4D). Subsequent GO enrichment analyses of differentially active kinases identified significant associations with essential biological processes, including protein metabolic process, phosphorylation regulation, etc, and molecular functions encompassing serine/threonine kinase catalytic activity (Figure 4E). Through systematic interrogation of the DGIdb 5.0 pharmacogenomic database [25], we identified 28 *S. japonicum* kinases (including AKT3, CDKs, MAPKs and others) exhibiting>60% sequence identity with human drug-targetable kinases (Tables 3 and S8). Meanwhile, 30 FDA-approved therapeutic agents are available for the homologous human kinases, which may be candidates for developing drugs against schistosomiasis (**Table 3**).

**Table 3.**
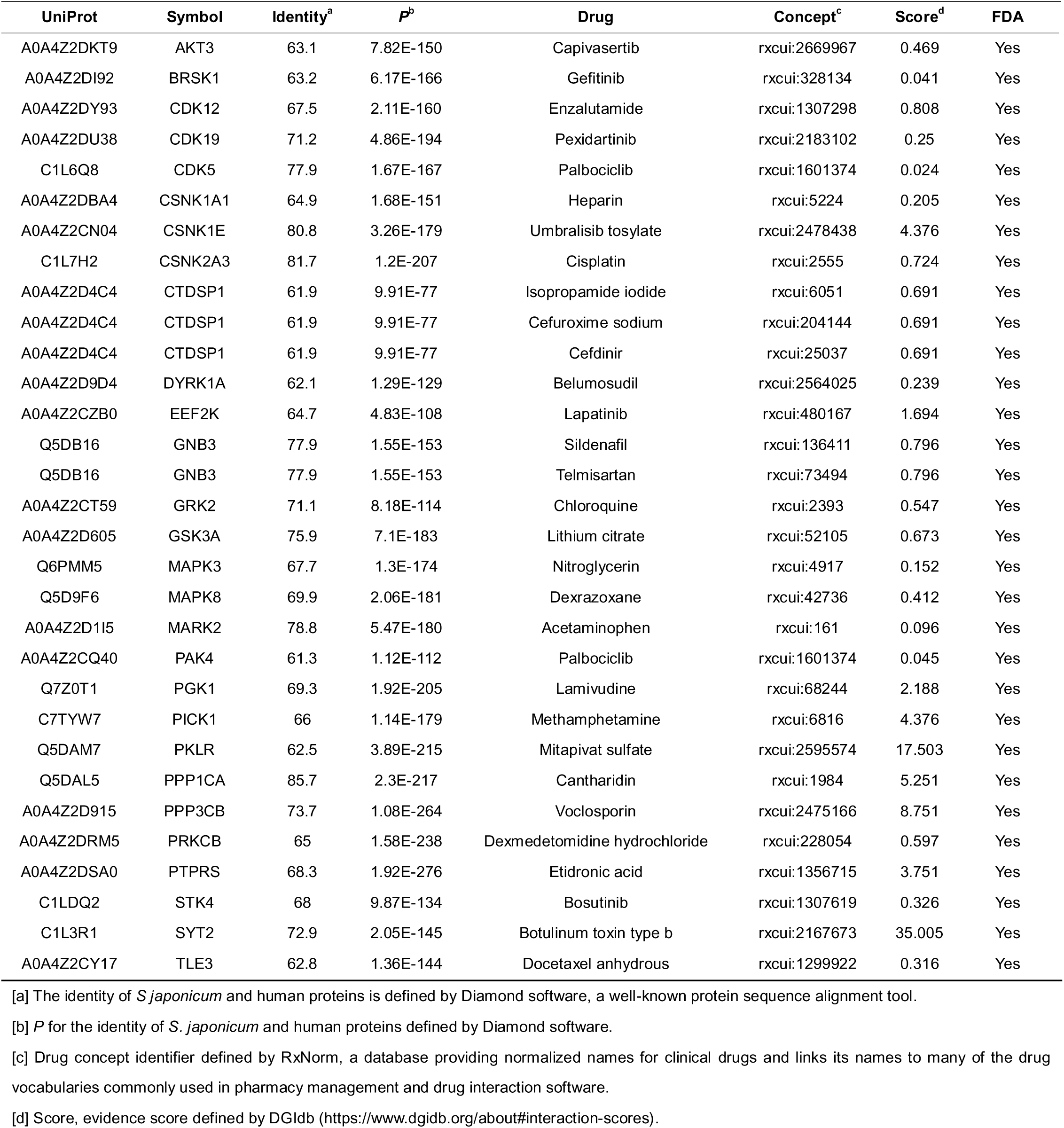
List of identified *Schistosoma japonicum* kinases with significant homology to human kinases targeted by FDA-approved drugs.

## Discussion

In recent years, multi-omics methodologies (e.g., proteomics and transcriptomics) have been systematically employed in schistosome research to advance gene annotation, spatiotemporal gene expression profiling, etc [12, 26, 27]. Notably, parasite eggs constitute the main pathogenic drivers of schistosomiasis pathogenesis [28, 29]. During sexual maturation and oviposition, male and female parasites fulfill distinct yet complementary biological roles, suggesting potential divergence in their protein ontologies and functional repertoires. To elucidate the molecular mechanisms underlying these processes, we conducted an iTRAQ-based quantitative proteomic and phosphoproteomic profiling to systematically characterize sex-biased protein and phosphoprotein profiles in adult *S. japonicum*.

Our analytical workflow successfully identified 10,046 non-phosphorylated peptides corresponding to 2,055 distinct proteins, establishing a comprehensive baseline proteomic inventory. In our study, cross-platform analyses revealed a considerable correlation between proteomic and transcriptomic profiles, and RNA levels can explain about 30% variance in protein levels. Except from transcriptional level regulation, protein abundance obtained in proteomic profiling can be explained at biological and technical layers, including: 1) post-transcriptional regulation (e.g., miRNA-mediated mRNA degradation or translational repression); 2) protein translation regulation (differences in ribosome binding efficiency or translation initiation rates that affect protein synthesis); 3) technical differences, such as different procedure of sample preparation (e.g., RNA vs. protein extraction methods), analyzed platforms (e.g., RNA-seq sensitivity vs. LC-MS/MS dynamic range), or data normalization strategies, etc [30, 31]. According to the literature, the protein variance that could be explained by mRNA variance varied quite in different study models, about 20%-80% [32]. Therefore, the findings in *S. japonicum* are consistent with the previous reports. Meanwhile, functional enrichment analyses demonstrated that highly abundant proteins were significantly associated with essential cellular processes, including cytoskeletal reorganization, microtubule dynamics, intercellular communication, and proteolytic regulation, corroborating and extending previous investigations in schistosome biology [12]. Actually, the dominant biological difference between female and male worms is coupled with their functional roles[33]. Female worms are primarily involved in egg production, which encompasses metabolic processes such as protein catabolism and proteolysis to generate vitellocytes. In contrast, male worms are enriched with proteins associated with cytoskeletal organization and contractile fiber components, which provide the necessary power for pairing. Collectively, the proteomics dataset generated from this study can reflect the molecular basis relate to the biological processes of *S. japonicum*.

Furthermore, to gain insight into the mechanisms of vitellocyte development, we integrated scRNA sequencing data of *S. mansoni* with our identified DEPs and identified that more than 15 DEGs were female-enriched (especially vitellocyte-enriched) genes, which may play roles in molecular material biological synthesis, substance transport, BMP signaling, etc. We hypothesized that BMP signaling is the dominant molecular mechanism in vitelline development.

Functional validation through RNA interference targeting female-enriched genes *Ft, Catb1,* and *Bmp* revealed significant impairment of vitellocyte development and oviposition for female worms. *Ft* encodes a protein named ferritin in *S. japonicum*. In 2022, Zeng et al. found that the ability of egg production was significantly reduced after ferritin knockdown [34]. *Catb1* encodes a protein named cathepsin B-like cysteine proteinase isoform 1, mainly expressed in gut cells. Cathepsin, a cysteine protease localized within subcellular endosomal and lysosomal compartments, is involved in the turnover of intracellular and extracellular proteins to mediate processes including apoptosis, pyroptosis, ferroptosis, necroptosis, and autophagic cell death [35]. BMPs (Bone morphogenetic protein) belong to the TGF-β superfamily and play crucial roles in the formation and regeneration of skeletal and cartilage tissue [36]. Although its biological roles in schistosomes are unclear, *Bmp* is versatile in multiple biological processes, including cell differentiation and fate determination, intestinal homeostasis and development, etc [36, 37]. In our experiments, we observed that silencing *Bmp* in *S. japonicum* can effectively inhibit vitelline development and cell proliferation in the ovary of female worms, which is similar to *Ft* and *Catb1* knockdown models. These results demonstrate that BMP signaling is implicated in the molecular mechanism of vitelline development of female worms. However, whether BMP signaling cross-talks with *Ft*/*Catb1* ontologies still needs further investigation.

Next, we developed a computational framework for kinase activity quantification through quantitative phosphorylated substrate profiling by the following paradigm steps: 1) kinase annotation using homology-based prediction through protein sequence alignment, 2) construction of kinase-substrate interaction networks (or gene sets) through the integration of functional and physical protein-protein interaction data; 3) quantitative kinase activity inference via GSVA-based algorithm using phosphoproteomic profiles mapped against curated substrate gene sets. By bridging protein-protein interaction data (such as the STRING database) with phosphoproteomic data, our integrative approach provides a systematic strategy to overcome the inherent challenges of studying the kinome of non-model organisms with limited prior annotations. Collectively, these methodological advancements could accelerate therapeutic discovery by pinpointing kinase-mediated developmental vulnerabilities that could be targeted by selective inhibitors.

The pivotal roles of kinases in schistosomes have been studied to some extent, revealing conserved and parasite-specific kinases that regulate critical developmental pathways, particularly in signaling, growth, differentiation, and reproductive processes. For instance, protein kinases (PKs), including serine/threonine kinases (e.g., PLKs, MAPKs, etc) and protein tyrosine kinases (PTKs), are functionally indispensable for key biological processes such as gonad maturation, pairing-dependent sexual differentiation, and host-parasite interactions [38, 39]. Cellular tyrosine kinases (CTKs), such as SmTK3, are implicated in female gonad differentiation and signaling cascades [40, 41]. RNA interference and pharmacological inhibition experiments have validated kinases like PKC, ERK, and FAK as potential drug targets [42, 43]. In the current study, DGIdb 5.0, a pharmacogenomic database, was used in our screening step, which directly prioritized the 30 FDA-approved kinase inhibitors. Among them, Chloroquine, a 4-aminoquinoline derivative, has been extensively used as a frontline antimalarial agent for over seven decades, demonstrating efficacy against *Plasmodium falciparum* and *Plasmodium berghei* strains [44]. Its mechanism involves accumulation within the acidic digestive vacuoles of intraerythrocytic malaria parasites. Next, Lamivudine, a synthetic cytosine nucleoside analog, exhibits robust therapeutic efficacy and favorable tolerability profiles in patients with chronic hepatitis B virus infection [45, 46]. This compound functions as a viral DNA chain terminator through competitive inhibition of reverse transcriptase enzymatic activity. Furthermore, lamivudine interacts with the kinase domains, including those of EGFR, RIPK1, and RIPK3, implicating its role in the regulation of necroptosis and suggesting putative antineoplastic mechanisms [47]. In 2020, Melissa and coworkers demonstrated that lapatinib has potent bioactive properties against adult *S. mansoni* [48]. Investigations by Fatemeh and coworkers revealed that topical administration of 0.1% cantharidin over two weeks achieves marked therapeutic efficacy against cutaneous leishmaniasis in murine models [49]. Eman and coworkers elucidated cisplatin’s anti-schistosomal activity, evidenced by tegumental damage, reduced anti-schistosomal IgG levels, and decreased egg burden, among other effects [50]. Overall, these studies corroborated the credibility of our findings regarding our computational framework to identify effective inhibitors targeting *S. japonicum* kinases. However, there are also challenges regarding side effects when developing kinase inhibitors targeting parasitic lesions due to selectivity concerns of the leading candidates. Therefore, optimization and modification steps may improve to exploit structural differences in parasite kinase active sites [51], screen of FDA-approved kinase inhibitors against parasite-specific kinases [52], and focus on essential parasite-specific kinases with unique structural features [53, 54].

In summary, our study provides: 1) a quantitative atlas of sex-biased proteomes and phosphoproteomes for adult *S. japonicum*; 2) mechanistic insights into vitelline development of female worms; 3) a computational framework for kinase activity profiling. These findings may provide important clues for understanding vitelline development and oviposition in *S. japonicum*.

## Materials and Methods

### Parasite infection and sample collection

In the current study, New Zealand rabbits and *Oncomelania hapensis* were used to maintain the life cycles of *S. japonicum* (Anhui isolate), and animals were cultured at Tongji University and Shanghai Veterinary Research Institute of the Chinese Academy of Agricultural Sciences. For the infection via the abdomen skin, high-activity *S. japonicum* cercariae (around 1500) were infected in rabbits. Adult schistosome worms were collected from the hepatic portal vein of the infected animals at 35 days post-infection (dpi). Worms were briefly cultured for around 4 hrs (at 37□) to separate female and male worms, which were further manually collected for protein extraction. Meanwhile, the worms were also snap-frozen in liquid nitrogen for further use.

### Protein extraction

Protein samples were obtained following a previous protocol with brief revisions [55]. In brief, around 50-100 mg liquid nitrogen frozen or fresh (adult female/male schistosomes) worms were homogenized into protein lysate using 500 µL lysis buffer (containing 1 mM DTT, 4% SDS, 150 mM Tris-HCl, pH8.0). The obtained lysates were heated for 5 min at 100□, and followed by ultrasonic disruption using a general program of 10 times 10 s sonication with 15 s intervals. Subsequently, the lysates were heated for 5 min at 100□ and followed by a 10 min centrifuge (at 10,000 g) to remove tissue or cell debris. Finally, the BCA protein assay kit (Cat: 23225, Thermo Scientific) was used to determine the protein concentration of the obtained samples. The quantified protein samples were stored at -80□ for further use.

### Protein digestion and iTRAQ labeling

Protein digestion was carried out according to the FASP protocol by Wiśniewski et al [55]. Briefly, each protein sample (around 200 μg) was added 30 μl SDT buffer (containing 4% SDS, 150 mM Tris-HCl, pH 8.0, 100 mM DTT). Repeated ultrafiltration (Pall units, 10 kD) was performed to remove detergents and other small molecules (e.g., DTT) in protein samples, using UA buffer (containing 8M Urea, 150 mM Tris-HCl, pH 8.0). Then, 100 μl 0.05M iodoacetamide in UA buffer was added to the protein samples to block reduced cysteine residues, and the tubes were placed in the dark for 20 min. Filters pretreated using 100 μl UA 100 μl DS buffer (50 mM triethylammoniumbicarbonate at pH 8.5) were used to filter the protein samples, and the obtained suspensions were further digested at 37□, using 2 μg trypsin (Promega, in 40 μl DS buffer). The obtained peptide concentration was determined using a UV light spectrophotometer at 280 nm.

The peptide samples were further labeled using the 8-plex iTRAQ (Isobaric Tags for Relative and Absolute Quantitation) reagent following the manufacturer’s protocol (Cat: 4352135, Sigma). Briefly, 70 μl of ethanol-dissolved iTRAQ reagent was added to obtain peptide samples labeled as Male-R1-113, Male-R2-114, Male-R3-115, Female-R1-116, Female-R2-117, and Female-R3-118. All the samples were dried under vacuum for further use.

### Phosphopeptide enrichment using TiO_2_ beads

Phosphopeptide enrichment was performed on the obtained four-plex iTRAQ-labeled peptides using TiO_2_ beads following the previous protocol [56]. Briefly, around 500 µL commercial loading buffer was added to each tube of the trypsin-digested peptide mixture to fully dilute the samples, and then TiO_2_ beads were added to enrich phosphopeptides. After a 40-minute shake, the TiO_2_ beads in each tube were packed into a GELoader tip (Cat: Z317047 Eppendorf, Hamburg, Germany). The column was sequentially washed using 50 μl washing buffer I (30% ACN and 3% TFA) three times, and then 50 μl washing buffer II (80% ACN and 0.3% TFA) three times. Next, 50 μl NH_4_OH (pH 10.5) was used to elute the bound peptides. Finally, the vacuum freeze-dried lyophilized phosphopeptides were resolved in 0.1% formic acid for Liquid Chromatograph-tandem Mass Spectrometer (LC-MS/MS) analyses.

### LC-MS/MS analyses

The obtained phosphopeptide samples were subjected to LC-MS/MS analyses using an automated Easy-nLC 1000 apparatus interfaced with a Q-Exactive mass spectrometer (Cat:IQLAAEGAAPFALGMAZR, Thermo Fisher Scientific). Briefly, 5 μL of the phosphopeptide samples was sequentially amalgamated with 15 μL of 0.1% (v/v) trifluoroacetic acid, 10 μL of the solution blend, and the obtained samples were subjected to LC-MS/MS analyses. During this procedure, a pre-column (20 mm×100 μm, 5 μM-C18) and an analytical column (250 mm×75 μm, 3 μM-C18) were used (Thermo Fisher Scientific), with mobile phases A (containing 0.1% formic acid in water) and B (containing 0.1% formic acid in 84% ACN). The phosphopeptides were fractionated (at a flow rate of 250 nL/min) using the gradient mobile phases: 0-55 % mobile phase B from 0-220 min, 55-100 % mobile phase B from 220-228 min, and 100% mobile phase B from 228–240 min.

For LC-MS/MS analyses, peptides were scrutinized in positive ion mode. In the MS spectra acquirement, a data-dependent top 10 methodology was employed to dynamically select the most abundant precursor ions from the survey scan (300-1800 m/z) for HCD (higher energy collisional dissociation) fragmentation. The predictive Automatic Gain Control (pAGC) method was used to determine the target value. Dynamic exclusion duration was set as 40.0s. A resolution of 70,000 at m/z 200 was used for survey scans, and 17,500 at m/z 200 for HCD spectra. The normalized collision energy was set to 27eV, and the under-fill ratio (stipulating the minimum percentage of the target value likely to be attained at maximum fill time) was defined as 0.1%. Peptide recognition mode was set to enabled. Data-dependent mass spectra were acquired for 240 min. The full MS surveys were collated over a mass-to-charge ratio (m/z) range of 300–1,800, with a resolution of 70,000 at m/z 200. For LC-MS/MS, a resolution of 17,500 at m/z 200 was used, with an isolation window of 2 m/z.

### Protein identification and data analysis

All label-free quantitative raw data files for proteome and phosphoproteome analyses were searched against a database based on two combined databases derived from the *S. japonicum* protein database (LSBI-Sjr, 12657 sequences; 4929382 residues) (http://lifecenter.sgst.cn/schistosoma/) and *S. japonicum* NCBI (51,526 entries) [57-59]. In detail, the raw files were analyzed individually with Mascot 2.2 with the following settings: 1) peptide mass tolerance = 20 ppm; 2) MS/MS tolerance = 0.05 Da; 3) Enzyme = Trypsin; 4) Missed cleavage = 2; 5) Fixed modification: Carbamidomethyl (C), iTRAQ4/8plex (K), iTRAQ4/8plex (N-term); 6) Variable modification: Oxidation (M); Phosphorylation (S/T/Y); 7) False discovery rates (FDR) lower than 1% at both the peptide and protein levels was set as an acceptance criterion for identifications. In the quantification using Proteome Discoverer 1.3, the peak intensity of each expected iTRAQ reporter ion was analyzed using the following parameters: 1) unique peptides with false discovery rate (FDR) ≤ 0.01; 2) excluding all quantification values without quantification channels, normalize on protein median, normalize all peptide ratios by the median protein ratio; and 3) median protein ratio = 1 after normalization [59]. For each phosphorylation site on the phosphopeptides, the probability of phosphorylation site was set to>75%, and the phosphorylation site score was> 50 [60]. Blast2GO was used to predict the biological processes of *S. japonicum* phosphoproteins [61]. STRING (http://string-db.org/) database was used to predict protein-protein interaction, which was visualized by the R package igraph (https://github.com/igraph/igraph). In the present study, the peptide FDR is lower than 1% for database searching. The DEPs were defined using DEqMS (R packages, a tool to perform statistical analyses of differential protein expression for quantitative proteomics data) [62], with a cutoff of adjusted *p<*0.05 and fold-change>1.5.

### RNA sequencing analyses

Samples utilized in LC-MS/MS analyses underwent total RNA isolation using TRIzol™ Reagent (Invitrogen) in accordance with the manufacturer’s protocol. RNA quality was assessed via the RNA Integrity Number (RIN) employing an Agilent 2100 system (Agilent Technologies). RNA samples with RIN >8 were subsequently selected for transcriptomic library construction and RNA sequencing. RNA sequencing was conducted at BGI (Beijing Genomics Institute, Shenzhen, China) following the manufacturer’s instructions.

The raw data underwent quality control using Trim-galore (v0.6.10, https://github.com/FelixKrueger/TrimGalore) to eliminate low-quality reads, adaptors, and reads containing poly-N sequences (with parameters -quality 30, -stringency 3). Subsequently, the obtained clean reads were quantified to generate a gene expression matrix using the Salmon software (v1.10.2, https://salmon.readthedocs.io/en/latest/salmon.html). In the quantification, the transcriptome sequences annotated based on assembly ASM636876v1 (https://www.ebi.ac.uk/ena/browser/view/GCA_006368765.1) of *S. japonicum* were used to build an index for the command “*salmon quant*”. For differential expression gene (DEG) analyses, the DESeq2 R package (v1.48.2, https://bioconductor.org/packages/release/bioc/html/DESeq2.html) was employed with the default settings, and DEG was defined by a cutoff of adjusted p<0.01 and fold-change>2.

### Single-cell RNA sequencing analyses

Single-cell RNA (scRNA) sequencing data of *S. mansomi* by Wendt et al. were downloaded from NCBI Gene Expression Omnibus under accession number GSE146737 [17]. Seurat package (https://satijalab.org/seurat/), a widely adopted framework in bioinformatics, was employed to analyze the scRNA sequencing data and the visualization, following the official analytical workflow. We adopted the cell type annotation in the published paper.

### Real-time qPCR (RT-qPCR)

Total RNAs were extracted from adult male and female worms (35 dpi) using Trizol (Thermo Fisher Scientific) and real-time RT-PCR was carried out following the manufacturer’s protocol, Briefly, around 1 μg total RNA was subjected to a reverse transcription reaction using multi-Scribe reverse transcriptase reagent containing random hexamers (Eppendorf), incubated at 25□ for 10 min, 48□ for 30 min, and 95L for 5 min. RT-qPCR was carried out using 1 μl cDNA in a final volume of 25 μl containing 12.5 μl 2×One-Step TB Green PrimeScript RT-PCR Kit (Cat: RR066A, Takara), 10.5 μl H_2_O, and 1 μl (10 μM) specific primers for the corresponding genes on a Mastercycler ep realplex (Eppendorf). The following thermal cycling profile was used for the RT-qPCR reaction: 95□ for 30 s, followed by 40 cycles of amplification (95□ for 5s, 62□ for 30 s). The primers involved in the study are listed in Table S7.

For normalization, the primers of *nicotinamide adenine dinucleotide dehydrogenase* (*NADH*) (forward primer: CGA GGA CCT AAC AGC AGA GG; reverse primer: TCC GAA CGA ACT TTG AAT CC) were used as an internal control. In the present study, the transcript of target genes relative to *NADH* was quantified using the following algorithm: N = 2^−ΔCt^, ΔCt = Ct^target^ − Ct^NADH^ [63].

### dsRNA preparation and treatment for schistosomes

According to the manufacturer’s protocol, long dsRNA was synthesized *in vitro* using the T7 Megascript RNAi Kit (Cat: AM1333, Thermo Fisher Scientific), using PCR products (as the template). The *in vitro* transcription reactions were performed at 37□ for 12 hrs, followed by DNase treatment to remove the template. Finally, the recovered dsRNA products were run on a 1% agarose gel to check their integrity. The obtained dsRNAs described above were added to the culture medium at a concentration of 30 µg/ml on days 1, 2, and 5, respectively. Finally, mRNA suppression levels were measured via RT-qPCR at 6 days post-treatment.

### RNA interference using siRNA

Three siRNA duplexes targeting EWB00-001858 duplexes (siRNA-392, siRNA-211, siRNA-824), three siRNA duplexes targeting EWB00-010322 (siRNA-265, siRNA-1612, siRNA-2465), and one negative control siRNA (Supplementary Table 7) were designed, chemically synthesized, and annealed in Shanghai GenePharma (China).

Each dsRNA duplex (3 μg per shot) was electroporated (125 V, 20 ms, 1 pulse in 200 μl RPMI 1640 media) into cultured worms (25 dpi). Then, the worms were transferred into a 12-well cell culture plate containing 2 □mL complete RPMI-1640 media supplemented with 2 □g/L glucose, 2.0□ g/L NaHCO_3_, 0.3 Lg/L L-glutamine, 15% fetal bovine serum (heat-inactivated), and 5% pen/strep (10□mg/streptomycin and 10,000 units penicillin in 0.9%□NaCl). At 48, 96, or 108 hrs of post-electroporation, the worms were collected for total RNA isolation in the RT-qPCR analyses. To address potential electroporation-induced effects, all groups (both control and RNAi treatment worms) were subjected to electroporation. Worm viability was rigorously assessed before motility analyses, and only intact, actively moving worms were selected for measurements.

### FastBlue BB staining

For FastBlue BB staining, worms from both the interference group and the control group were collected. The obtained worms were fixed in 4% formaldehyde for 4 hrs at room temperature, then washed with PBSTx with shaking for 10 min at room temperature. The freshly prepared 1% FastBlue BB staining solution was filtered through a 0.22 μm filter. After staining the parasites for 3 min, the parasites were washed with PBSTx for 3×5 min. Subsequently, the worms were washed using 80% ethanol (for 10 min). Finally, the mounted slides were observed under an optical microscope to visualize the staining results.

### Worm motility

After electroporated with inhibitors, male and female worms (25 dpi) were cultured for 5 days *in vitro*. Morphologic alterations and motility of worms (10 worms/well) were daily monitored using an inverted microscope.

### Statistical analyses

GraphPad Prism 9 (Boston, USA) software was used for statistical analyses and visualization. Student’s *t*-test or ANOVA was employed for comparing the means of 2 or >2 groups, respectively. The difference with *p<*0.05 is considered statistically significance.

## Ethics statement

All experiments involving mice were carried out based on the Guide for the Care and Use of Laboratory Animals of the Ministry of Science and Technology of the People’s Republic of China. All efforts were made to minimize animal suffering. All animal procedures were approved by the Animal Management Committee and the Animal Care and Use Committee of the Shanghai Science and Technology Commission of the Shanghai municipal government for Tongji University (TJAA00822501).

## Data availability

The MS raw datasets were submitted to the National Genomics Data Center, Beijing Institute of Genomics, Chinese Academy of Sciences / China National Center for Bioinformatics under the project PRJCA040825.

## Credit authorship contribution statement

**Chuantao Fang**: Conceptualization, Investigation, Software, Formal analyses, Visualization, Writing-original draft. **Bikash Giri**: Formal analyses, Investigation, Visualization, Writing-original draft. **Guofeng Cheng**: Conceptualization, Investigation, Writing-original draft, Writing-review & editing. All authors have read and approved the final manuscript.

## Acknowledgments

This study was supported by the National Natural Science Foundation of China (Grant No.: 31472187) and the Fundamental Research Funds for the Central Universities from Tongji University of China (Grant No.: 22120220010). The study was also sponsored by the Shanghai Tongji University Education Development Foundation of China.

## Declaration of AI and AI-assisted technologies in the writing process

We did not use any AI tools to assist the process of manuscript writing.

## Supplementary material

**Figure S1 Overview of iBQP and iBQPP protein profiles**

**A.** Functional enrichment analyses of highly expressed proteins in proteomics using GO and KEGG gene sets. MF, molecular function of the GO database; KEGG, Kyoto Encyclopedia of Genes and Genomes. The top 20 enriched terms were presented.

**Figure S2 Differentially expressed proteins between male and female worms.**

**A**. Principal component analyses of proteomic profile dataset. **B**. Boxplot showed the total count of peptides among the samples. **C**. The correlation of mRNAs (by RNA-seq) and proteins (by proteomics) abundance of *S. japonicum.* The mRNA levels were quantified using TPM (Transcripts Per Kilobase of exon model per Million mapped reads), and the protein levels were defined by the normalized peptide count using peptide length. **D**. Heatmap of differentially expressed genes between male and female worms. The cutoff of adjusted p < 0.01 and fold-change > 2 was used. **E**. Volcano plot showing the differentially expressed genes/proteins between male and female worms. **F**. Dotplot showing the DEG expression pattern in different cell types of a scRNA sequencing dataset of schistosomes. **G**. Protein-protein interaction of identified differentially expressed proteins. The STRINGdb R package was employed for the analyses.

**Figure S3 Differentially expressed proteins between male and female worms.**

**A**. Functional enrichment analyses for protein interactors in Figure S2G using gene sets defined by the Gene Ontology and KEGG. BP, biological processes; CC, cellular components; MF, molecular function. p*<*0.05 was used as a cutoff. The top 10 enriched terms were presented. **B.** Real-time PCR reveals the knockdown efficiency of siRNA targeting the indicated protein-coding genes. Independent siRNA targeting different sites was designed. *** p <0.001. **C.** Worm mobility showed the functional alteration of worms after knocking down sex-specific coding genes. ***p <0.001 and ns, no significance.

**Figure S4 Differentially expressed kinases between male and female.**

**A**. Boxplot showed the total count of phosphorylated peptides among the samples. **B**. Distribution of unique peptide coverage/total peptide of 1,833 phosphorylated proteins identified. **C**. The distribution of peptides on full-length protein (Top) and peptide length (Bottom). **D**. The motif of specific kinases by the Motifx software.

**Figure S5 Specific kinase motif by MEME**

The motifs of the indicated kinases are predicted by MEME software. Default settings were used in the analyses.

**Supplementary Table S1. The basic information of iTRAQ-based quantitative proteome**

**Supplementary Table S2. Expression matrix of iTRAQ-based quantitative proteome**

**Supplementary Table S3. Differentially expressed proteins of iTRAQ-based quantitative proteome**

**Supplementary Table S4. Differentially expressed genes between males and females from the public RNA-seq dataset of *S. japonicum***

**Supplementary Table S5. The basic information of iTRAQ-based quantitative phosphoproteome**

**Supplementary Table S6. Differentially expressed phosphorylated proteins of iTRAQ-based quantitative proteome**

**Supplementary Table S7. siRNA sequence for targeting the indicated genes and primer sequences used in the study**

**Supplementary Table S8. The list of drugs for potentially targeting *S. japonicum* kinases**

