## Supplementary figures and images for "Quantitative proteomics and phosphoproteomics analyses identify sex-biased protein ontologies of *Schistosoma japonicum*"

### Supplemental Fig 1-5

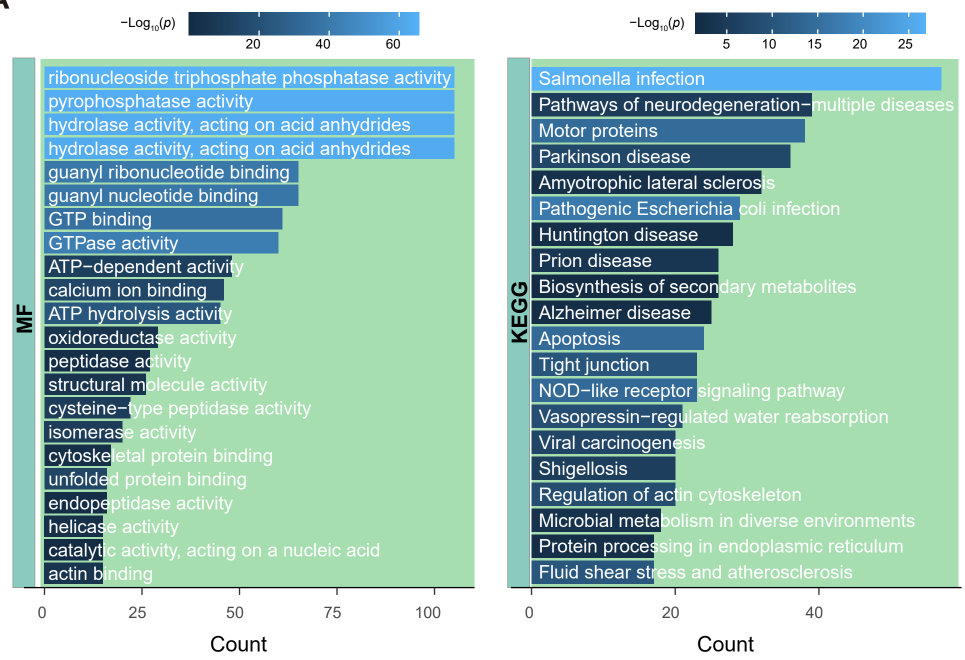

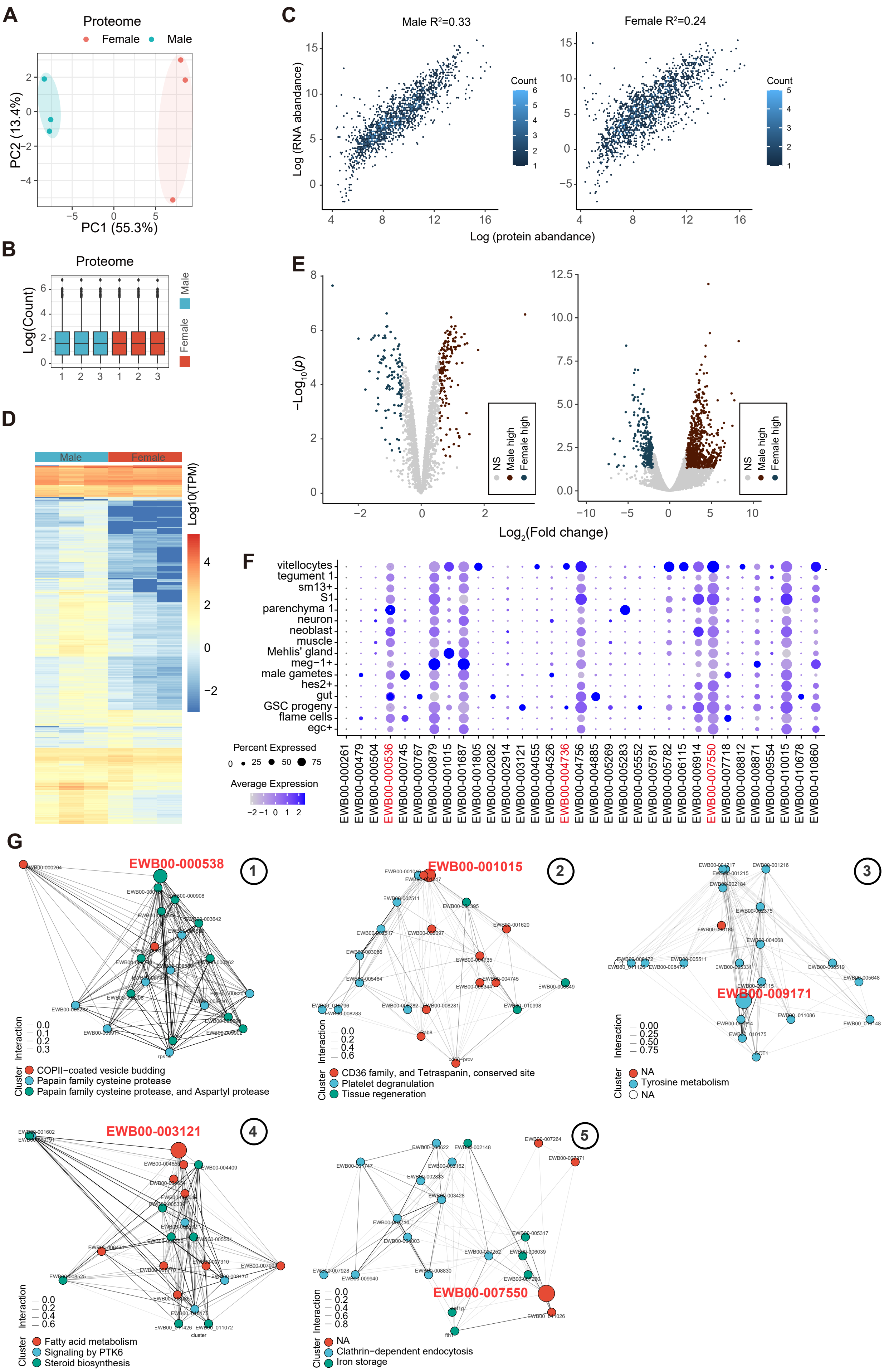

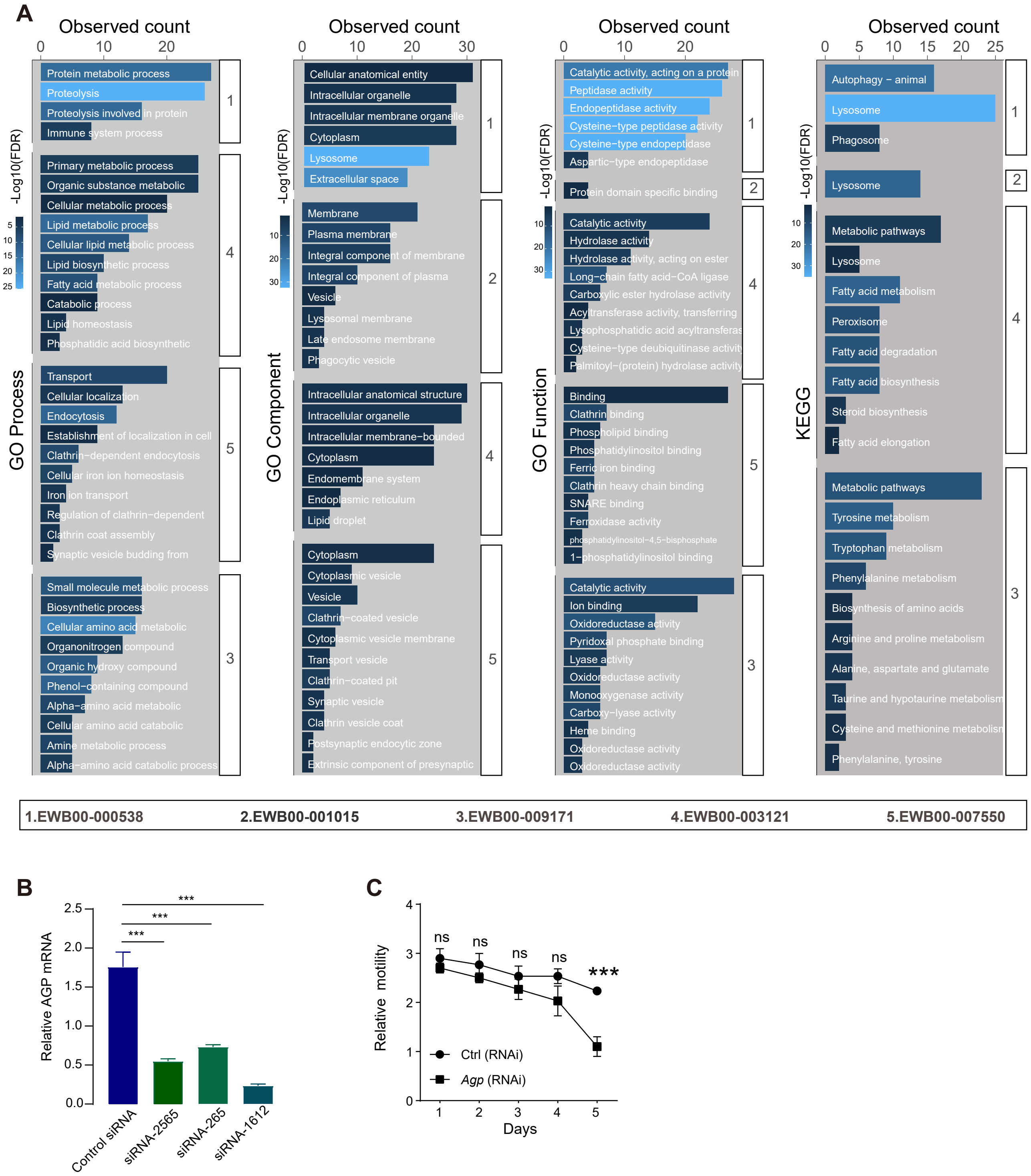

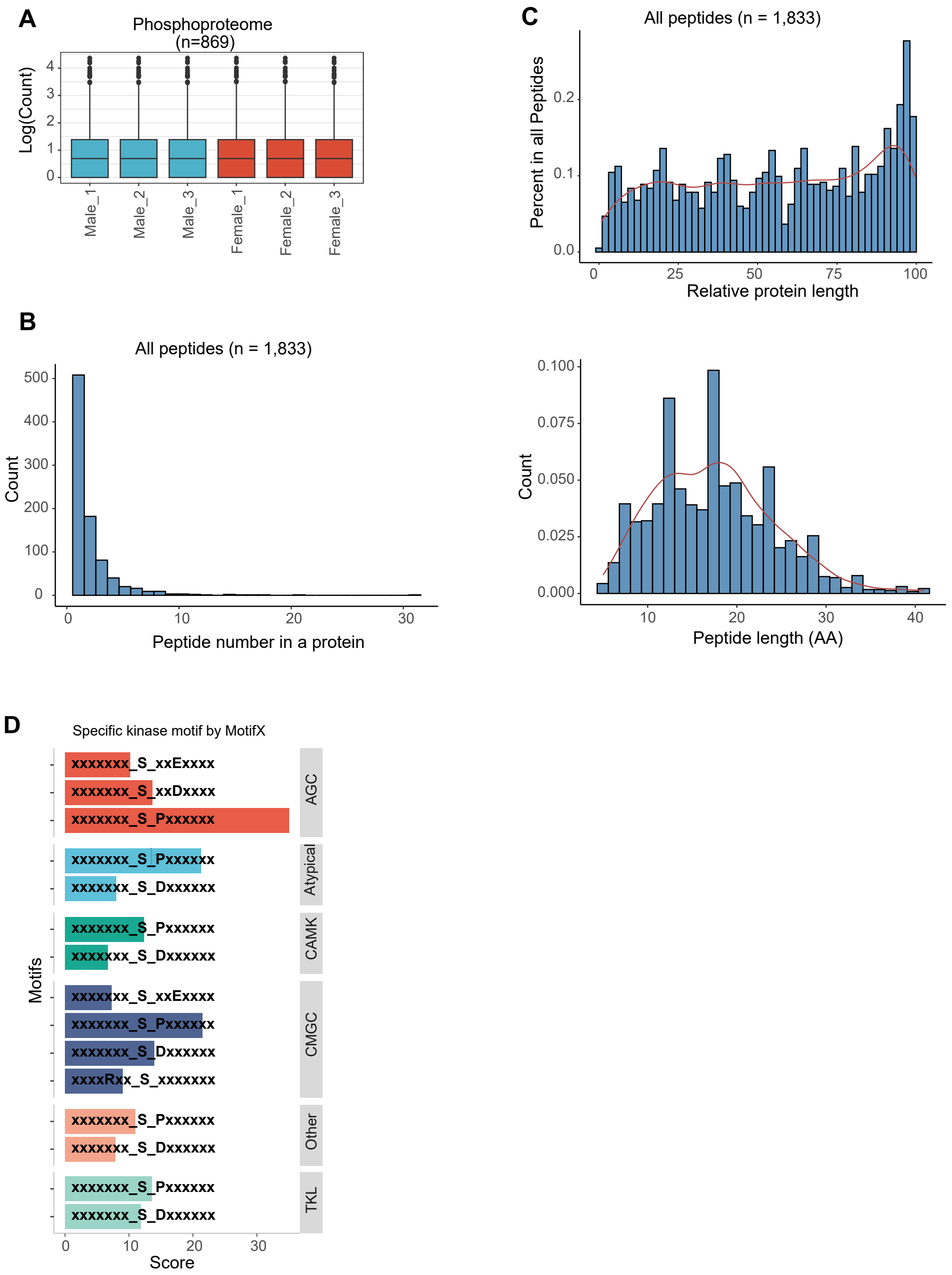

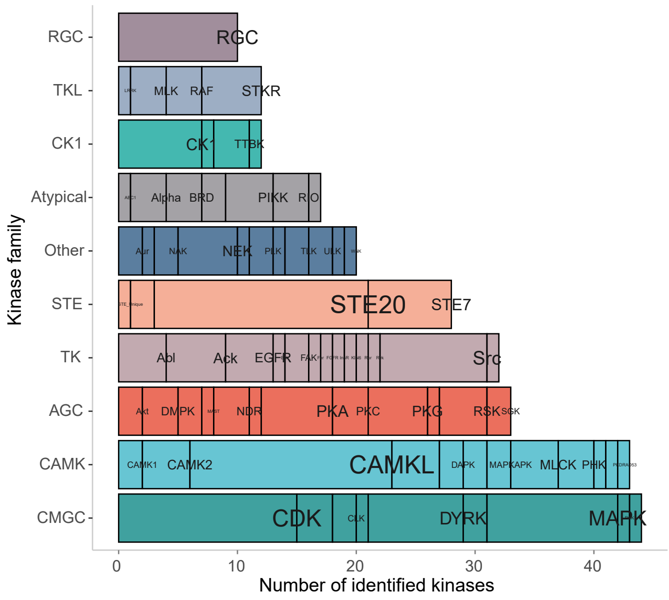
